# Designing BH3-mimetic Peptide Inhibitors for the Viral Bcl-2 Homologs A179L and BHRF1: Importance of long-range electrostatic interactions

**DOI:** 10.1101/2021.06.23.449612

**Authors:** C. Narendra Reddy, Ramasubbu Sankararamakrishnan

## Abstract

Viruses have evolved strategies to prevent apoptosis of infected cells at early stages of infection. The viral proteins (vBcl-2s) from specific viral genes adopt a helical fold that is structurally similar to that of mammalian anti-apoptotic Bcl-2 proteins and exhibit little sequence similarity. Hence vBcl-2 homologs are attractive targets to prevent viral infection. However, very few studies have focused on developing inhibitors for vBcl-2 homologs. In this study, we have considered two vBcl-2 homologs, A179L from African swine fever virus and BHRF1 from Epstein-Barr virus. We generated two sets of 8000 randomized BH3-like sequences from eight wild-type pro-apoptotic BH3 peptides. During this process, the four conserved hydrophobic residues and an Asp residue were retained at their respective positions and all other positions were substituted randomly without any bias. We constructed 8000 structures each for A179L and BHRF1 in complex with BH3-like sequences. Histograms of interaction energies calculated between the peptide and the protein resulted in negatively skewed distributions. The BH3-like peptides with high helical propensities selected from the negative tail of respective interaction energy distributions exhibited more favorable interactions with A179L and BHRF1 and they are rich in basic residues. Molecular dynamics studies and electrostatic potential maps further revealed that both acidic and basic residues favorably interact with A179L while only basic residues have the most favorable interactions with BHRF1. As in mammalian homologs, the role of long range interactions and non-hotspot residues have to be taken into account while designing specific BH3-mimetic inhibitors for vBcl-2 homologs.

## Introduction

The immediate response of viral infection in mammals includes programmed cell death (PCD) of infected cells and immune response. Among the many different types of cell deaths, apoptosis is an important mode of PCD in mammalian cells and can occur either via extrinsic or intrinsic pathways ^1^. Intrinsic pathway or mitochondrial apoptosis is mediated by Bcl-2 family of proteins and the family members include those that promote (pro-apoptotic) or oppose (anti-apoptotic) cell death ^2–3^. Anti-apoptotic Bcl-2 members are characterized by the presence of multi Bcl-2 homology (BH) domains (BH1 to BH4) while pro-apoptotic members have either multi BH domains (BH1 to BH3) or only BH3 domain ^4–5^. “BH3-only” pro-apoptotic proteins can be activators of pro-apoptotic Bak/Bax or can be repressors antagonizing the anti-apoptotic Bcl-2 proteins such as Bcl-X_L_ or Mcl-1 ^6–7^ Upon activation, Bax/Bak accumulate on the mitochondrial surface and form pores in mitochondrial outer membranes. This process enables apoptogenic proteins like cytochrome C to be released in cytoplasm leading to eventual cell death ^8–9^. At the initial stages of infection, viruses have evolved mechanisms to inhibit apoptosis to avoid early cell death that will help the virus to replicate ^10–12^ This is achieved through viral proteins that mimic the mammalian anti-apoptotic Bcl-2 proteins. Many viral Bcl-2 homologs (vBcl-2) are structurally similar to their mammalian counterparts with distinct Bcl-2 α-helical fold ^13–16^. However, they exhibit little or no sequence similarity with the anti-apoptotic Bcl-2 proteins. Examples include F1L and N1 from vaccinia virus (PDB ID: 2VTY and 2UXE) ^17–18^, M11L from myxoma virus (PDB ID: 2O42) ^19^, BHRF1 from Epstein-Barr virus (EBV) (PDB ID: 2WH6) ^20^, KsBcl-2 from Kaposi sarcoma herpesvirus (PDB ID: 1K3K)^21^, F1L from Variola virus (PDB ID: 5AJJ) ^22^, M11 from γ-herpesvirus 68 (PDB ID: 2ABO) ^23^, ORFV125 from ORF virus (PDB ID: 7ADS) ^24^, GIV66 from grouper iridovirus, (PDB ID: 5VMN) ^25^, CNP058 from canarypox virus (PDB ID: 5WOS) ^26^, FPV039 from fowlpox virus (PDB ID: 5TZP) ^27^, 16L protein from tanapoxvirus (PDB ID: 6TQP) ^28^ and A179L from African swine fever virus (PDB ID: 5UA4)^29^. These viruses cause diseases such as canarypox (song birds), smallpox (humans), myxomatosis (rabbits), Burkitt’s lymphoma, Hodgkin and non-Hodgkin lymphoma and other cancers (humans), Castleman’s disease (humans), contagious ecthyma (sheep and goats), sleepy grouper disease (fish), fowlpox (chicken) and African swine fever (pigs and warthogs) ^30–37^. Thus the viral diseases can be lethal to humans and/or economically devastating. Hence, understanding the molecular mechanism of viral infection at the early stages of viral entry into host cells is an important step in curtailing the viral diseases.

In mammalian anti-apoptotic Bcl-2 proteins, it has been shown that the peptide corresponding to the BH3-domain of pro-apoptotic counterparts is enough to trigger the apoptotic process ^38–39^. Several anti-apoptotic Bcl-2 proteins exhibit different affinities for pro-apoptotic BH3 peptides ^40–41^. While Bim binds to all anti-apoptotic Bcl-2 proteins, Bad binds to only Bcl-X_L_, Bcl-2 and Bcl-w and does not bind to Mcl-1 and A1. Similarly, Noxa binds to Mcl-1 and A1 but has no affinity for Bcl-X_L_, Bcl-w and Bcl-2. Binding studies for viral Bcl-2 homologs reveal that each vBcl-2 homolog has its own binding profile and many times they differ from mammalian pro-survival proteins^14, 17, 20, 22, 27, 29, 42–45^. Binding profiles of vBcl-2 homolog from African swine fever virus A179L show that it is a panproapoptotic Bcl-2 binder, binding to all BH3 motifs of various prodeath Bcl-2 proteins tested ^29^. On the other hand, F1L from vaccinia virus binds only to a limited number of pro-apoptotic BH3 peptides^17^.

Structural studies have shown that the pro-apoptotic BH3 peptides bind to the hydrophobic groove formed by many amphipathic α-helices of pro-survival Bcl-2 proteins^15–16,38^. Efforts to design small molecules that mimic the pro-apoptotic BH3 domain have been successful to some extent and some of them have reached clinical trials ^46–47^. However, these molecules are designed to antagonize mammalian anti-apoptotic Bcl-2 proteins. As vBcl-2s adopt a similar structural fold, one can imagine that the inhibitors developed for anti-apoptotic Bcl-2 proteins can also act on vBcl2s. Robert et al. tested ABT-737, a small molecule that specifically inhibits Bcl-2, Bcl-X_L_ and Bcl-w, on EBV-induced lymphoproliferative disorders like Burkitt’s lymphoma ^48^. This drug, alone or in combination with other conventional drugs, did not show any improvement on the overall survival of Burkitt’s lymphoma xenograft mice. Burrer et al. identified BH3-mimetic peptide that can selectively bind vBcl-2 from Kaposi’s sarcoma-associated herpesvirus by screening BH3 peptide libraries ^49^ However, the identified peptide could not cross the plasma membrane. There are few reports available describing the design of specific inhibitors for inactivating viral Bcl-2 homologs ^50^. Recently, David Baker and his colleagues designed a novel protein that can bind to BHRF1 of EBV with high affinity and their structural studies showed that this protein binds to the hydrophobic groove of BHRF1 and makes many additional contacts outside this region also ^51^. They also demonstrated that this protein triggers apoptosis in EBV-infected cell lines. In mouse xenograft disease model, the designed protein could suppress tumor progression and extended the lifespan of infected mice. Although these studies show promise in developing inhibitors specific to vBcl-2, these are too few compared to the numerous studies reported in the literature describing the design and development of inhibitors for mammalian anti-apoptotic Bcl-2 proteins. Hence, it is important that specific focus should be given in developing vBcl-2-specific inhibitors that can be used in immunocompromised infected patients.

We have recently developed a computational method to design BH3-like peptides that can bind specifically to either Bcl-X_L_ or Mcl-1^52^. Our studies showed that BH3-mimetic peptides have distinct preference for charged residues when they bind to Mcl-1 or Bcl-X_L_ and also highlighted the role of long range electrostatic interactions due to non-hotspot residues. The computationally designed BH3 peptide inhibitors, predicted to bind Mcl-1 or Bcl-XL with high affinity, were successfully validated in cell viability and cell proliferation studies. In the current study, we have applied the same approach to design BH3-like peptides for two of the vBcl-2 homologs, BHRF1 from Epstein-Barr virus and A179L from African swine fever virus. A179L binds to all pro-apoptotic BH3 peptides ^29^ while BHRF binds to a subset of them with high affinity including Bak and Bim ^20^. Results of this study show that the designed BH3-mimetic peptides are rich in basic residues for both BHRF1 and A179L. However, the pattern of interactions between the charged residues and the protein seems to differ between BHRF1 and A179L.

## Results

Wild-type BH3 sequences from pro-apoptotic Bcl-2 proteins were used to generate randomized BH3-like sequences (Figure 1) as described in the Methods section. We considered a total of eight BH3 wild-type sequences from Bak, Bad, Bid, Bik, Bim, Bmf, Noxa and Puma and in each case we generated two sets of 1000 BH3-like sequences. Structures of BH3-like sequences in complex with A179L were modeled as described in the Methods Section. The same exercise was repeated for generating BHRF1:BH3-like sequence complex structures. In total, we modeled 16,000 complex structures (8000 each for A179L and BHRF1 systems). Interaction energies between the peptide and protein (A179L or BHRF1) were calculated (see Methods section). Histograms of interaction energies of peptide: protein for BH3-like sequences derived from each wild-type BH3 peptide were plotted and are shown in Figure 2 and Figure 3 for A179L and BHRF1 respectively. It is evident that the interaction energies show negatively skewed extreme value distributions. We made a similar observation for mammalian anti-apoptotic Bcl-2 proteins also ^52^. In majority of the cases, the interaction energy of the wild-type peptide lies close to or within one standard deviation from the mean value. The BH3-like sequences whose interaction energies lie in the extreme negative tail region have the most favorable interactions with the protein. Hence, our interest is to select BH3-like sequences from this region which are predicted to have higher helicity and mean hydrophobic moment. It has been shown that helical stability of BH3 peptides is directly correlated with the binding affinity and the BH3-domain forms amphipathic helices in naturally occurring pro-apoptotic proteins^53–55^. For each wild-type BH3 peptide, we selected five BH3-like sequences from the extreme negative tail region of histograms with high predicted helicity. Helicity and mean helical hydrophobic moment of each BH3-peptide was predicted using Agadir and HELIQUEST web servers ^56–59^ We analyzed interaction energies of these selected peptides with the protein. The van der Waals and columbic components of interaction energies calculated for BH3-like sequences are compared with the corresponding wild-type pro-apoptotic BH3 peptide from which these sequences were derived. The following sections will first discuss the findings independently for A179L and BHRF1 and then the results for these proteins will be compared between them and also with respect to the mammalian anti-apoptotic proteins Bcl-X_L_ and Mcl-1.

**Figure 1:**
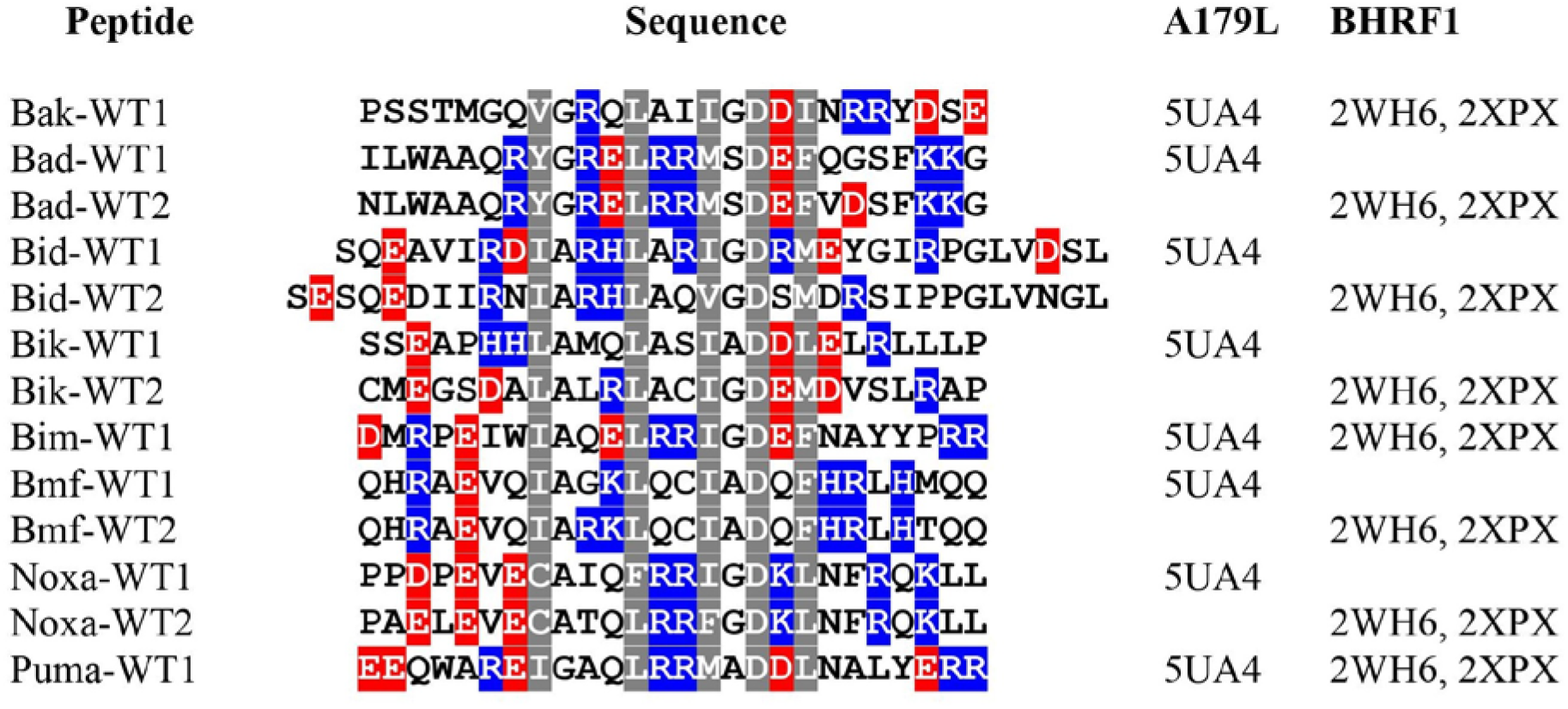
Sequences of wild-type of BH3 peptides from pro-apoptotic proteins that are used to derive BH3-like sequences. The residues which are retained while deriving BH3-like sequences are shown in gray background. Residues shown in red and blue background represent acidic and basic residues. PDB IDs of template structures which are used to model A179L and BHRF1 complex structures are provided.

**Figure 2:**
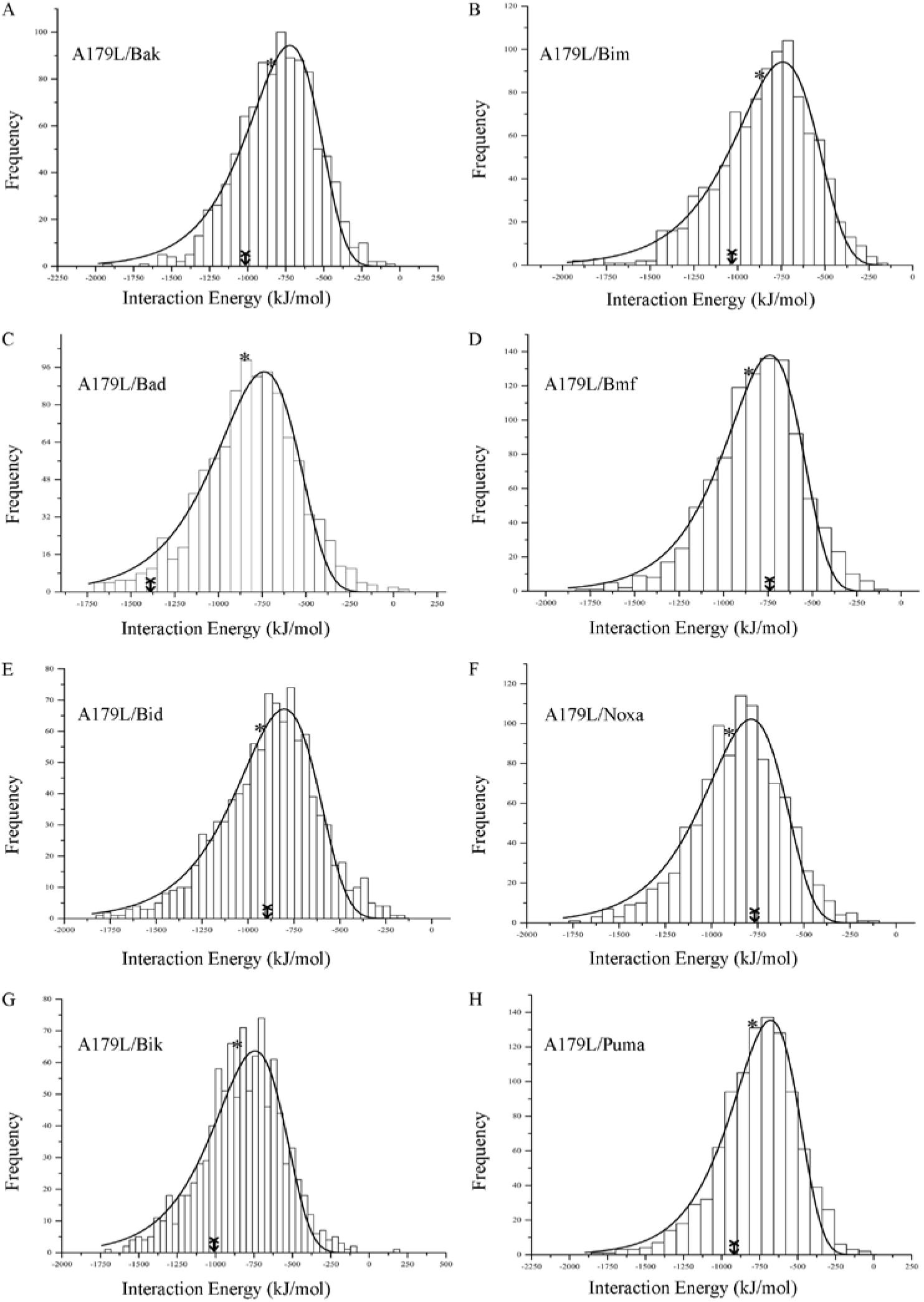
Histograms of interaction energies between A179L and BH3-like sequences. The wildtype BH3 peptides from which the BH3-like sequences were derived are indicated for each histogram. In each histogram, the mean value and the interaction energy of the wild-type BH3 peptide are designated with an asterisk and arrow respectively.

**Figure 3:**
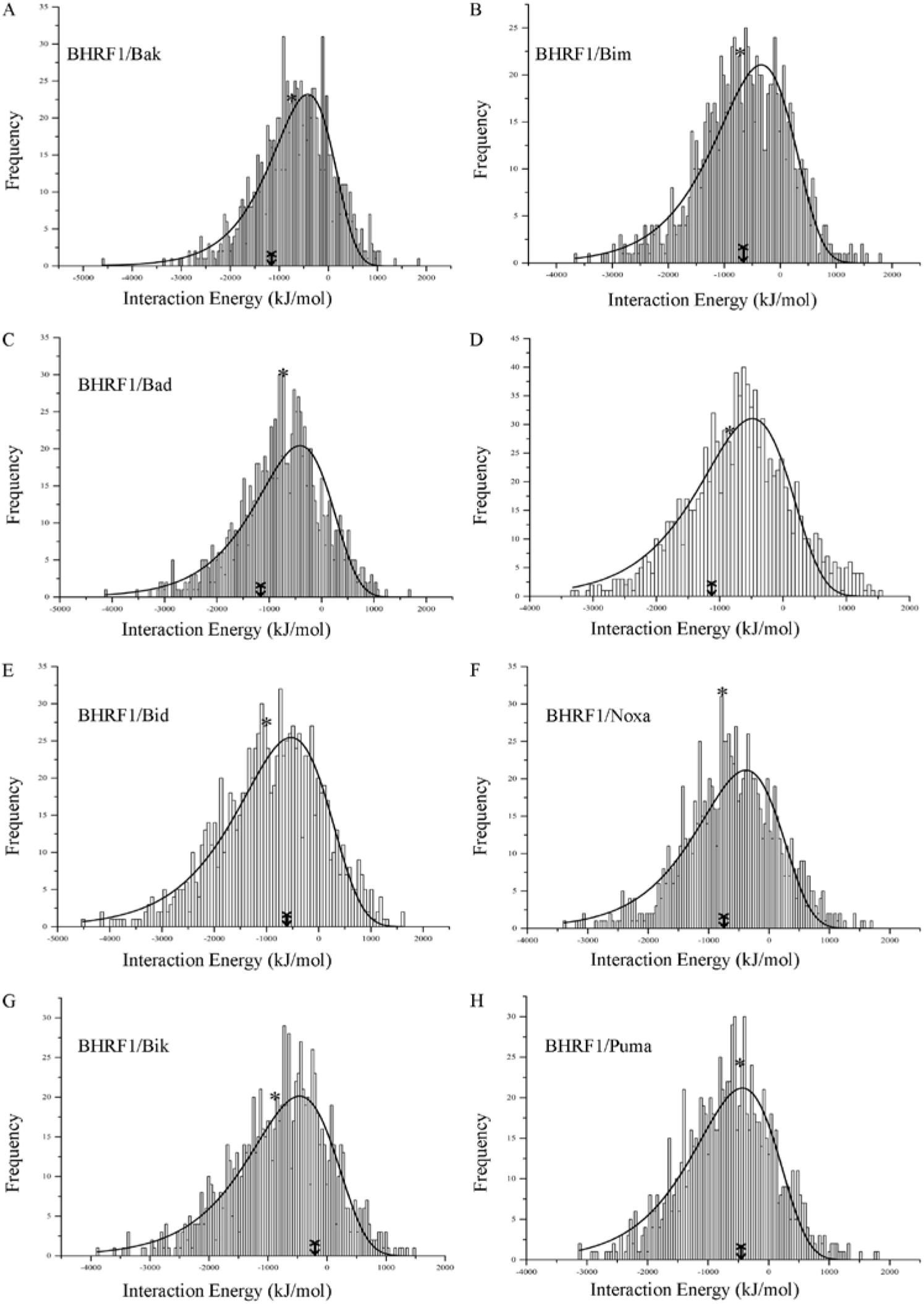
Histograms of interaction energies between BHRF1 and BH3-like sequences. The wild-type BH3 peptides from which the BH3-like sequences were derived are labeled in each histogram. The mean value and the interaction energy of the wild-type BH3 peptide in each histogram are designated with an asterisk and arrow respectively.

### A179L in complex with BH3-like sequences

We have selected top five BH3-like peptides derived from each of the eight wild-type pro-apoptotic BH3-sequences. These top BH3-like peptides were selected from the negative tail region of interaction energy distributions obtained for A179L modeled complex structures (Figure 2). The sequences of these peptides are shown in Figure 4 and their predicted helical propensities (Figure S1A) are significantly higher than that of corresponding wild-type BH3 peptides. The mean helical hydrophobic moment calculated for each BH3-like sequence is comparable to that of wild-type BH3 sequence from which they are derived (Figure S2A). The five selected BH3-like sequences derived from Bak are designated as Bak-1 to Bak-5. A similar notation was used for BH3-like sequences derived from other pro-apoptotic wild-type BH3 peptides. The selected sequences clearly show a pattern and they are rich in basic residues. The net charge of majority of these peptides varies from +1 to +9. Interaction energies of each of the selected BH3-like peptide in complex with A179L are plotted in Figure 5A. It is very clear that the BH3-like sequences interact with A179L much more favorably than the corresponding wildtype BH3 peptides. Interaction energies calculated for the wild-type peptides vary from −740 kJ/mol (Bmf-WT) to −1388 kJ/mol (Bad-WT). The BH3-like sequences derived from the wildtype BH3 peptides interact with A179L with interaction energies ranging from −1321 kJ/mol to − 1814 kJ/mol. In all the cases, the BH3-like sequences interact with A179L more favorably by 300 to 900 kJ/mol. Only Bad-3 interacts with A179L with similar interaction energy as that of wild-type Bad. When we examined the histogram of interaction energies of BH3-like sequences derived from Bad, we observe that the Bad wild-type peptide’s interaction energy with A179L lies near the negative tail of the distribution indicating that Bad wild-type sequence is already optimized to interact with A179L with most favorable interaction energy. The columbic component of interaction energies (Figure 5B) significantly contribute to the total interaction energies while van der Waals interaction energies between the selected peptides and A179L are comparable to the wild-type (Figure S3A). This indicates that the electrostatic interactions due to charged residues in the peptides play a major role in giving rise to highly favorable interaction energies with A179L.

**Figure 4:**
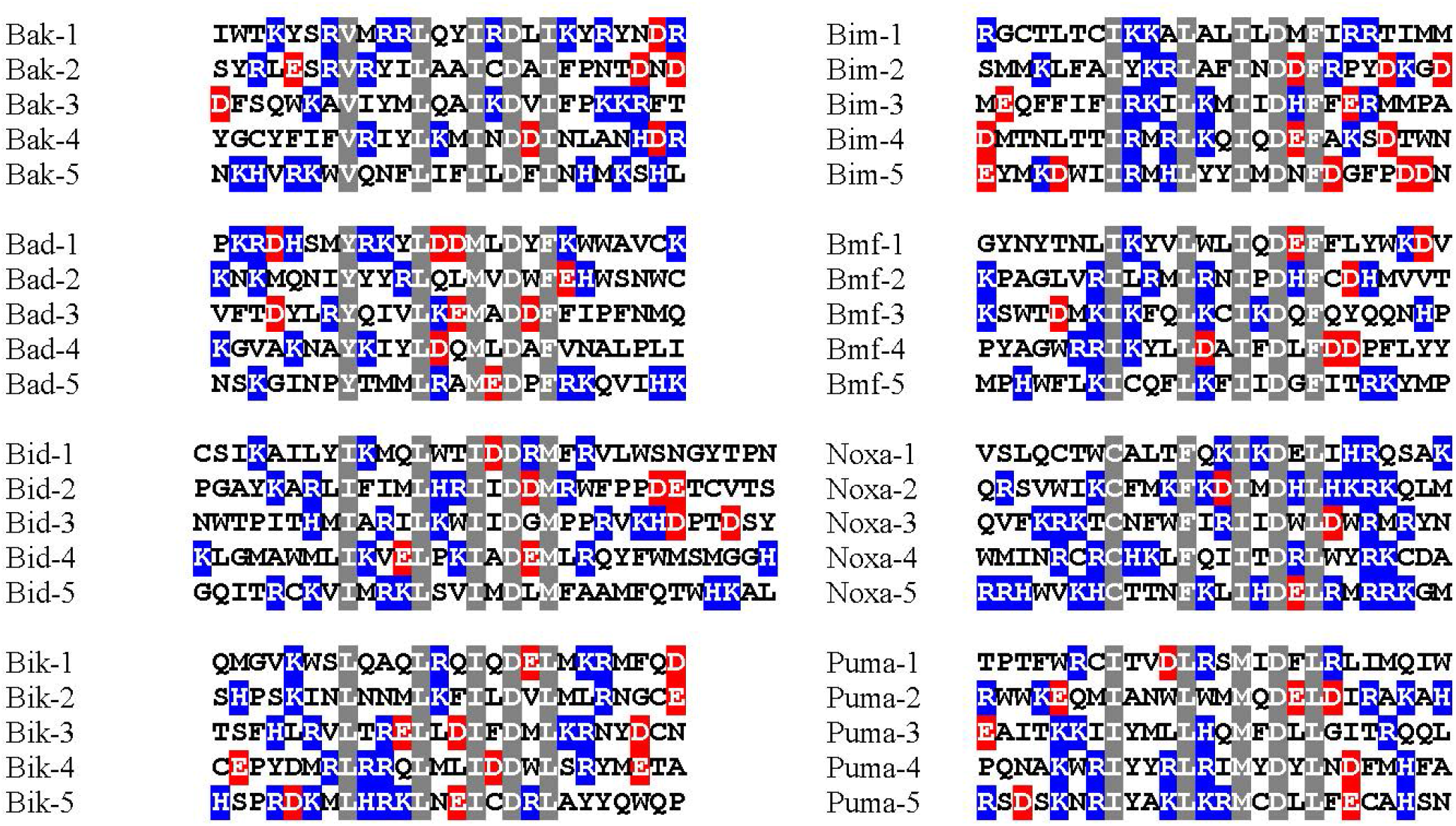
Top BH3-like sequences with high helical propensities and most favorable interaction energies with A179L selected from the extreme negative tail regions of respective interaction energy histograms (see Figure 2). The acidic, basic and conserved residues are shown in red, blue and gray background respectively.

**Figure 5:**
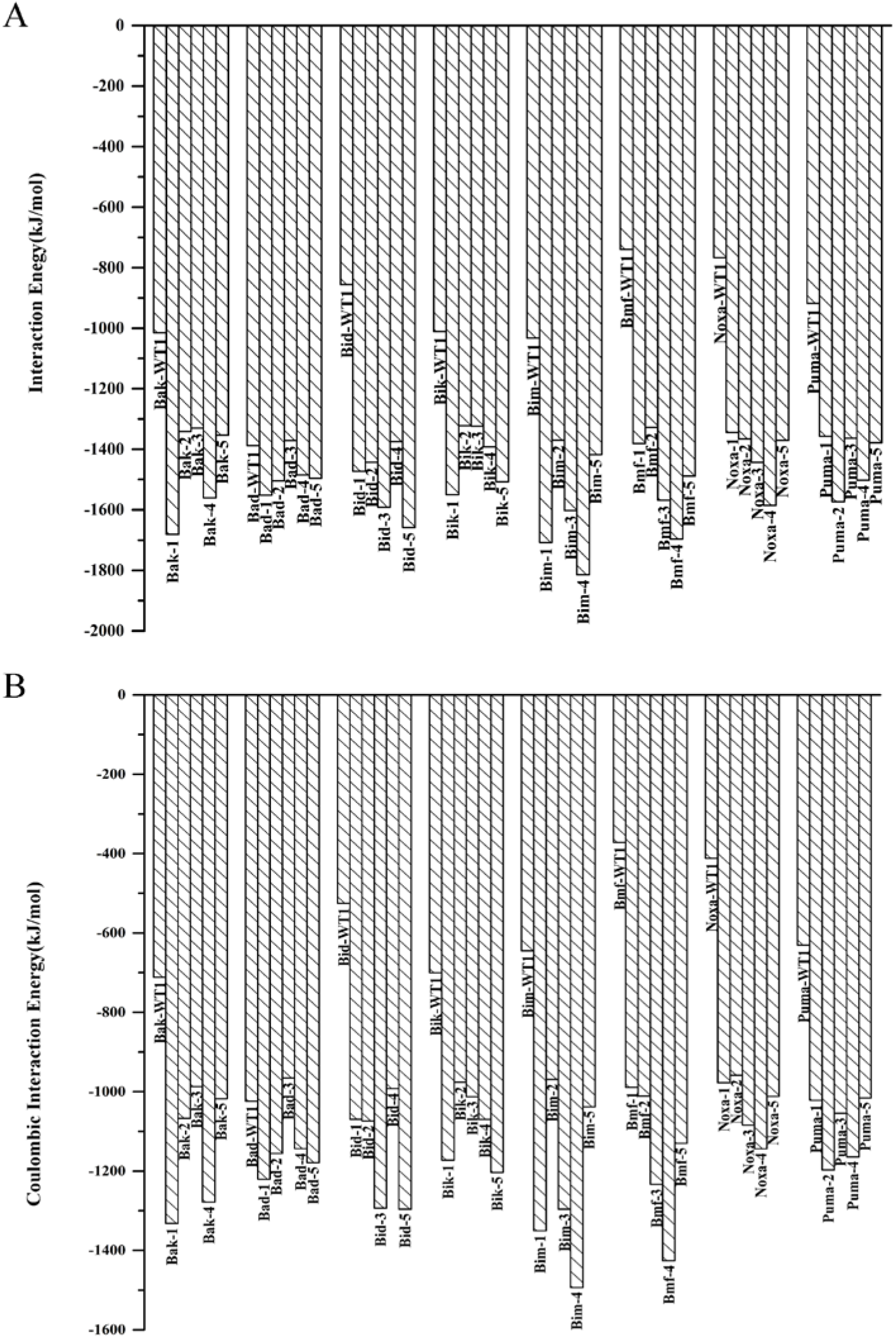
(A) Interaction energies of wild-type BH3 peptides and top selected BH3-like peptides in complex with A179L. (B) Electrostatic components of interaction energies of BH3 wild-type peptides and BH3-like sequences when bound to A179L.

### BHRF1:BH3-like peptide complex structures

The top five selected BH3-like sequences derived from each of the eight wild-type pro-apoptotic BH3 peptides are shown in Figure 6. These peptides were from the extreme negative tail of regions of interaction energy distributions for BHRF1 complex structures (Figure 3). Majority of the selected peptides have helical propensities significantly higher than that of wildtype BH3 peptides (Figure S1B). In most of the cases, the calculated mean hydrophobic moments are comparable to the corresponding wild-type BH3 peptides (Figure S2B). The five selected BH3-like peptides that are derived from wild-type Bak BH3 peptide are labeled as Bak-a to Bak-e. The same notation is followed for other BH3-like peptides also and this will help to distinguish from the BH3-like peptides that are used to model A179L complex structures. When we examined the BH3-like sequences derived from various wild-type pro-apoptotic BH3 regions, they are strikingly rich in basic residues. The net charge of the peptides varied from +1 to +7. Interaction energies of each set of BH3-like sequences derived from a specific pro-apoptotic BH3 domain clearly demonstrate that the wild-type pro-apoptotic BH3 peptides interact with BHRF1 at least 1000 kJ/mol less favorable than the BH3-like peptides (Figure 7A). When wildtype BH3 peptides interact with BHRF1, the interaction energies range from −206 (Bik-WT) to − 1170 (Bak-WT) kJ/mol. However, the BH3-like sequences derived from wild-type pro-apoptotic BH3 domains interact with BHRF1 with interaction energies varying from −1875 kJ/mol to −3830 kJ/mol. The differences in interaction energies are especially pronounced for the BH3-like peptides derived from Bik-WT and Puma-WT. It is important to note that the interaction energies of wild-type BH3 peptides lie close to the average values of their respective distributions in the histograms. For the Bik and Puma wild-type peptides, the interaction energies are more towards the positive side of the distributions. We have also calculated the columbic and van der Waals component of the interaction energies (Figure 7B and Figure S3B). It is evident that the major differences between the wild-type BH3 and BH3-like sequences are due to the electrostatic interactions. There is only marginal difference in the van der Waals component. It is especially important to note that the columbic component of interaction energies for Bik and Puma wild-type BH3 peptides is positive.

**Figure 6:**
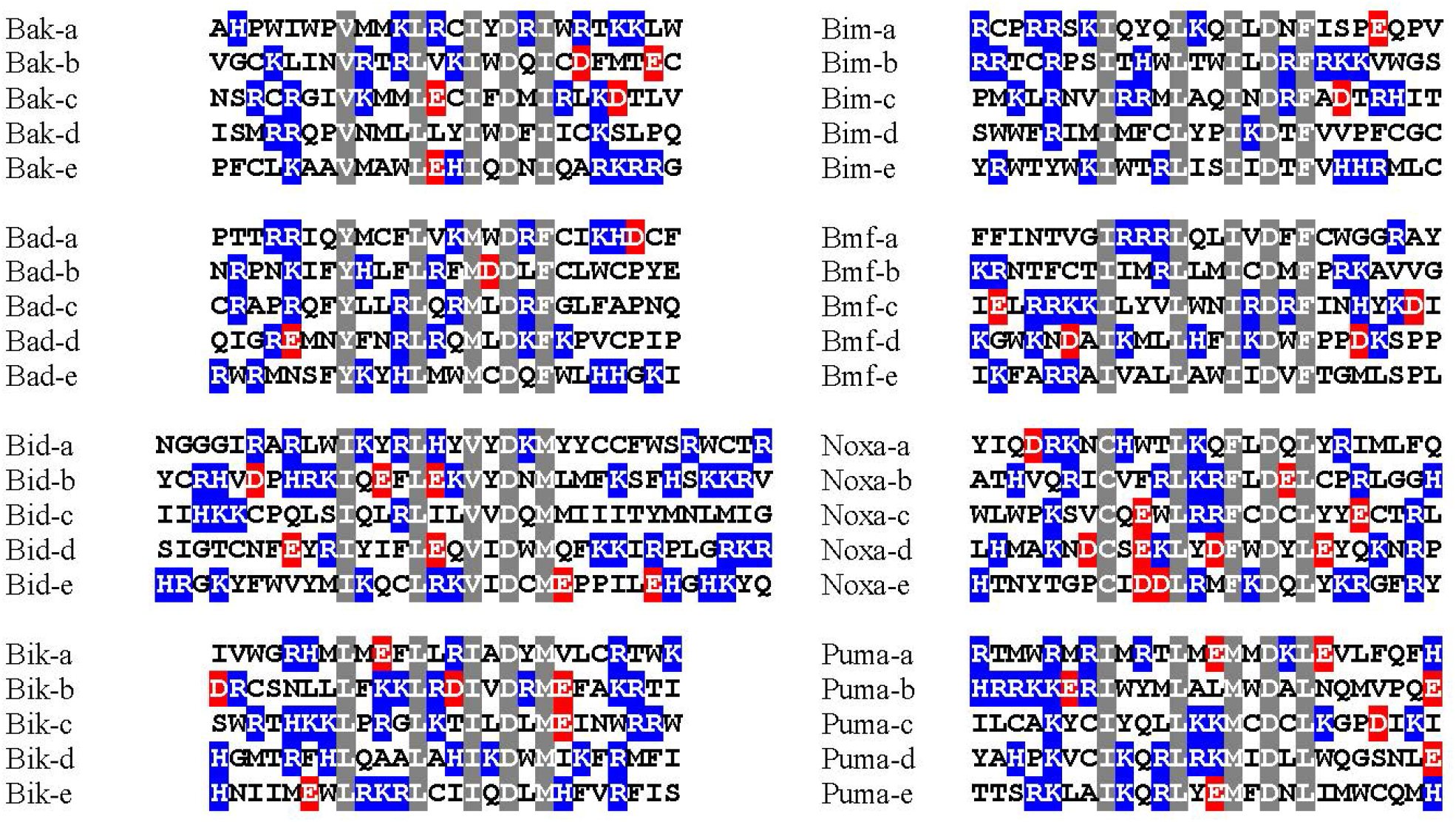
Top BH3-like peptides selected from the extreme negative tail end of interaction energy histograms (Figure 3). These peptides have high helical propensities and highly favorable interaction energies with BHRF1. The conserved hydrophobic residues and the conserved Asp are shown in gray background. The acidic and basic residues are displayed in red and blue background respectively.

**Figure 7:**
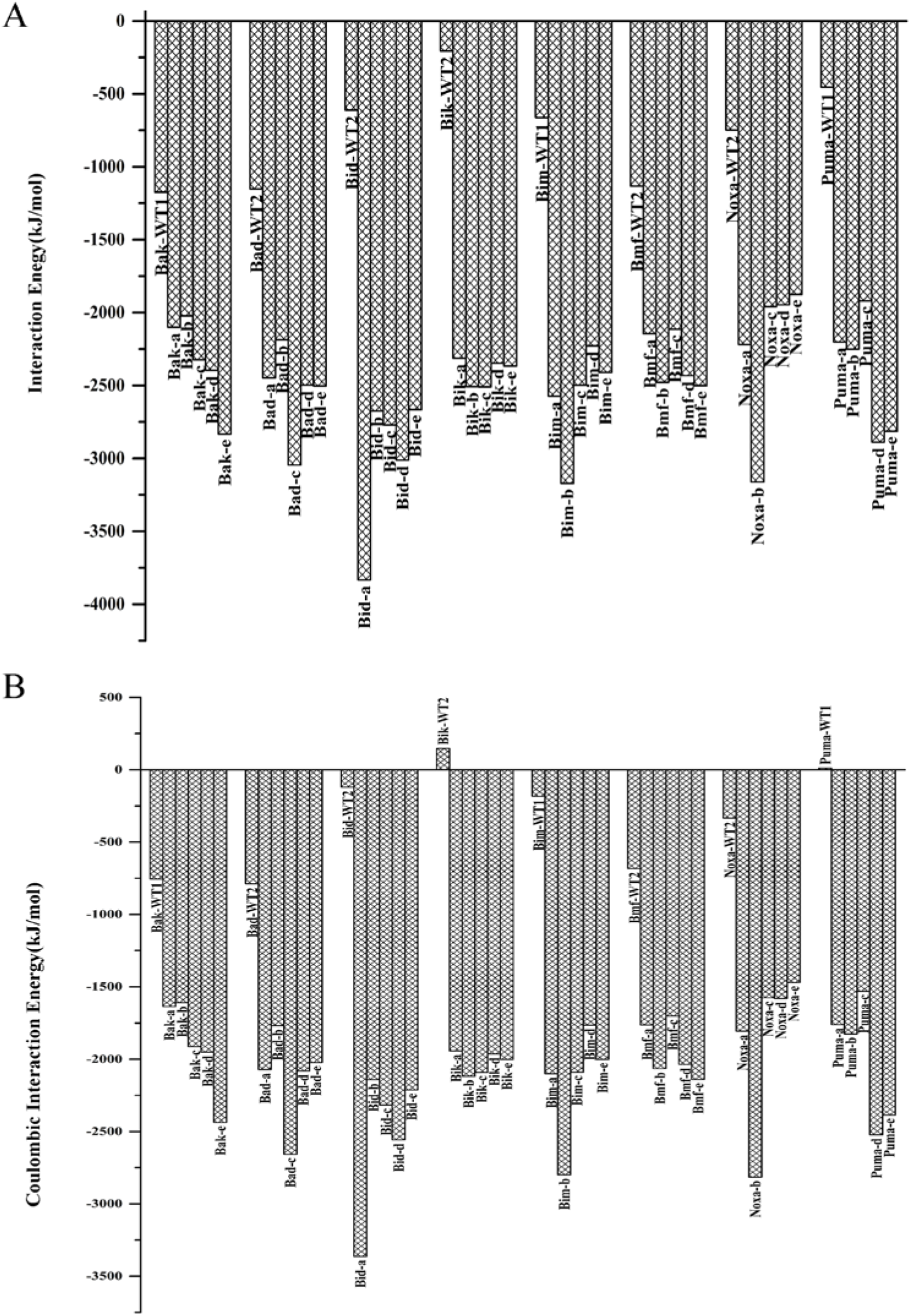
(A) Interaction energies of wild-type BH3 peptides and top selected BH3-like peptides in complex with BHRF1. (B) Electrostatic components of interaction energies of BH3 wild-type peptides and BH3-like sequences when bound to BHRF1.

### Molecular dynamics simulations of selected A179L and BHRF1 complex structures

Interaction energies calculated from the minimized complex structures gave an idea of the BH3-like peptides that prefer to bind to A179L or BHRF1 vBcl-2 homologs. However, it is possible that the interactions in the minimized structures may be transient and new interactions may form as protein-peptide complexes may evolve over a period of time. Hence, we carried out MD simulations on selected complex structures and compared the results with that of wild-type BH3 peptides in complex with A179L or BHRF1. MD simulations were performed as described in the Methods section. For A179L, we selected three BH3-like peptides (Bid-5, Bim-3 and Bmf-4). Similarly, we picked four BH3-like peptides in complex with BHRF1 (Bak-3, Bik-5, Bim-3 and Noxa-5). These peptides are specifically chosen because they have high predicted helical propensity and exhibit the most favorable interaction energies compared to their respective wildtype BH3 sequences. We performed 500 ns production run for each complex structure and compared the results with the A179L/BHRF1 structure in complex with pro-apoptotic wild-type BH3 peptides. A total of 14 simulations were carried out for a period of 7 p,s (Table S1). We have analyzed several properties including BH3 peptide helical stability and peptide:protein interaction energies.

DSSP plots of wild-type BH3 and BH3-like peptides bound to both A179L and BHRF1 show that the BH3 helical regions are mostly stable during the production runs (data not shown). The initial protein:peptide complex structure and the MD simulated complex structure saved at the end of 500 ns production run are shown in Figure S4 for each simulated system. It is evident that both BH3 wild-type peptides and the BH3-like peptides have mostly stable helical structures. Since the BH3-like sequences selected from the extreme negative tail regions of interaction energy distributions are rich in basic residues, we calculated the average interaction energies between the charged residues of the peptide and A179L/BHRF1 protein (Figure 8 and Figure 9) for all the simulated systems. When BH3-like peptides, Bmf-4 and Bid-5, and the wildtype BH3 peptides interact with A179L, both acidic and basic residues interact favorably with the protein (Figure 8). In the case of Bim-3, interaction energies of the acidic residues are slightly unfavorable. In general, the interaction energies of basic residues dominate giving rise to overall favorable interaction energies between the peptides and the protein. This also can be explained on the basis of electrostatic potential map of A179L (Figure 10B). The protein A179L has many acidic patches explaining why many basic residues are found in most favorably interacting BH3-like sequences. However, there are also regions of basic patch that can give favorable interactions from acidic residues. By interacting with complementary regions, both acidic and basic residues result in more favorable interaction energies and the charged residues, both positive and negative, seem to contribute to the highly favorable interaction energies as far as A179L is concerned (Figure 8).

**Figure 8:**
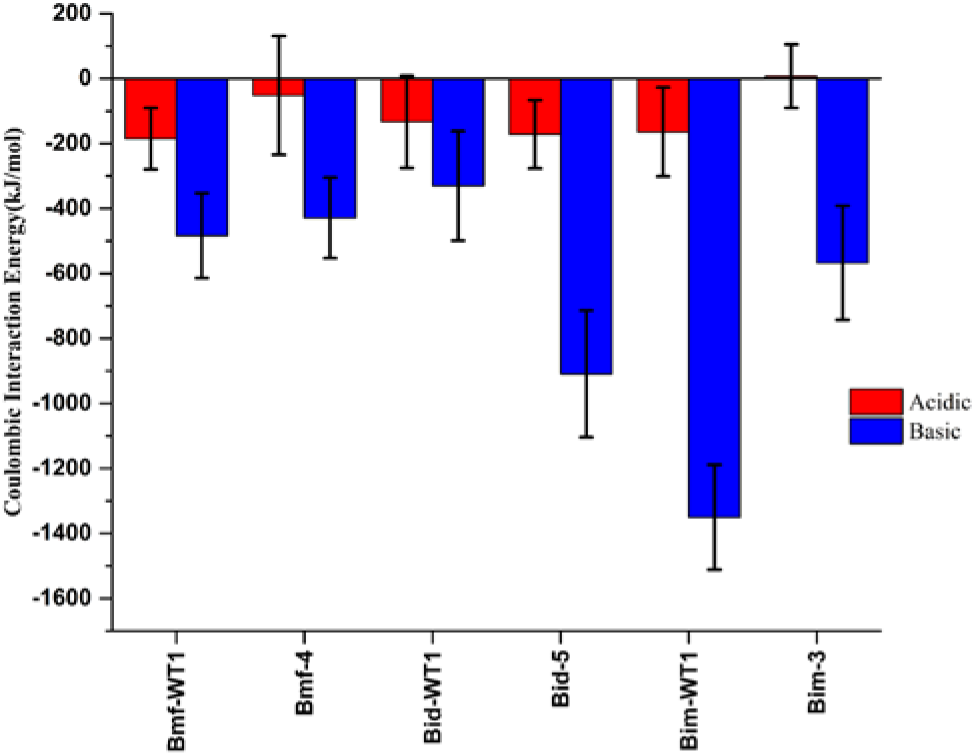
Average electrostatic interaction energies of A179L in complex with BH3-like sequences Bmf-4, Bid-5 and Bim-3 calculated from MD simulated structures. For comparison purpose, average electrostatic interaction energies of A179 in complex with the respective wildtype BH3 peptides are also shown. Electrostatic interactions calculated for acidic (red) and basic (blue) residues of the BH3 or BH3-like peptides are displayed separately.

**Figure 9:**
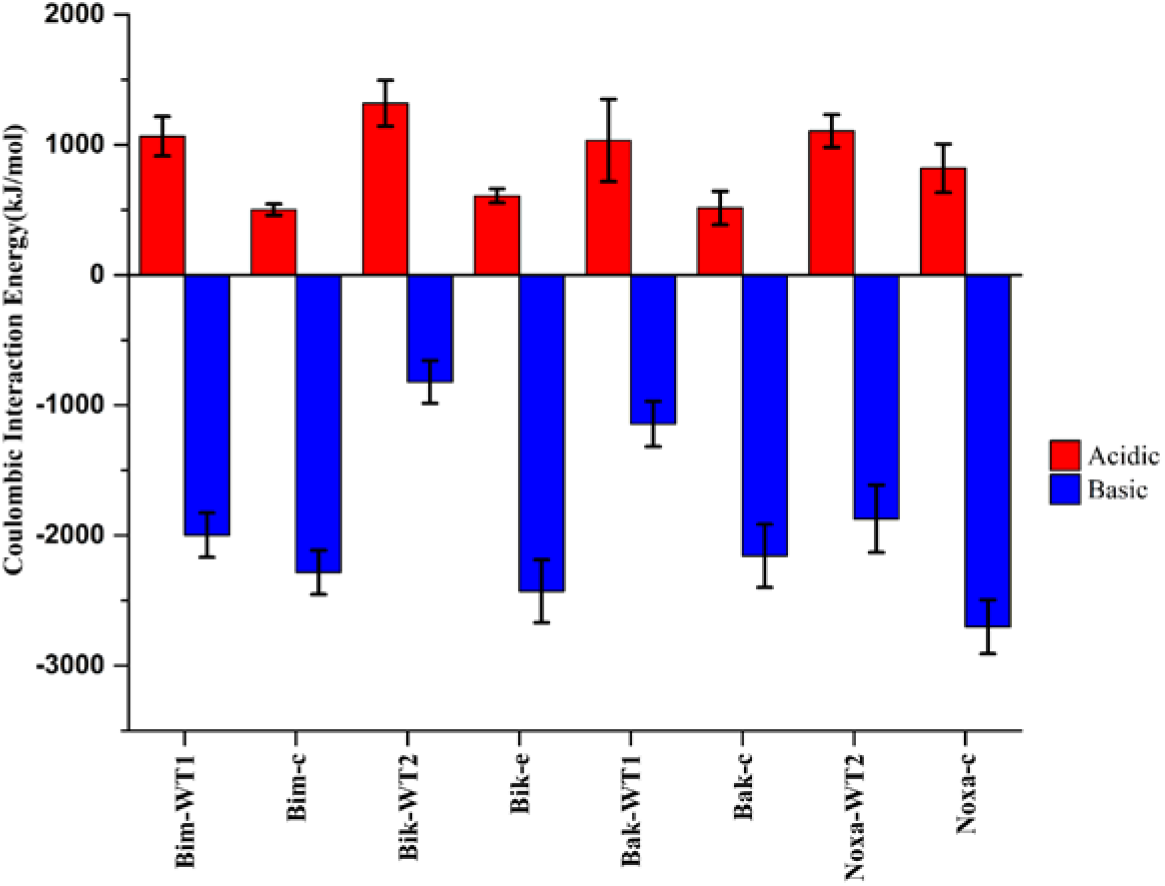
Average electrostatic interaction energies of BHRF1 in complex with BH3-like sequences Bim-c, Bik-e, Bak-c and Noxa-c calculated from MD simulated structures. For comparison purpose, average electrostatic interaction energies of A179 in complex with the respective wild-type BH3 peptides are also shown. Electrostatic interactions calculated for acidic (red) and basic (blue) residues of the BH3 or BH3-like peptides are displayed separately.

**Figure 10:**
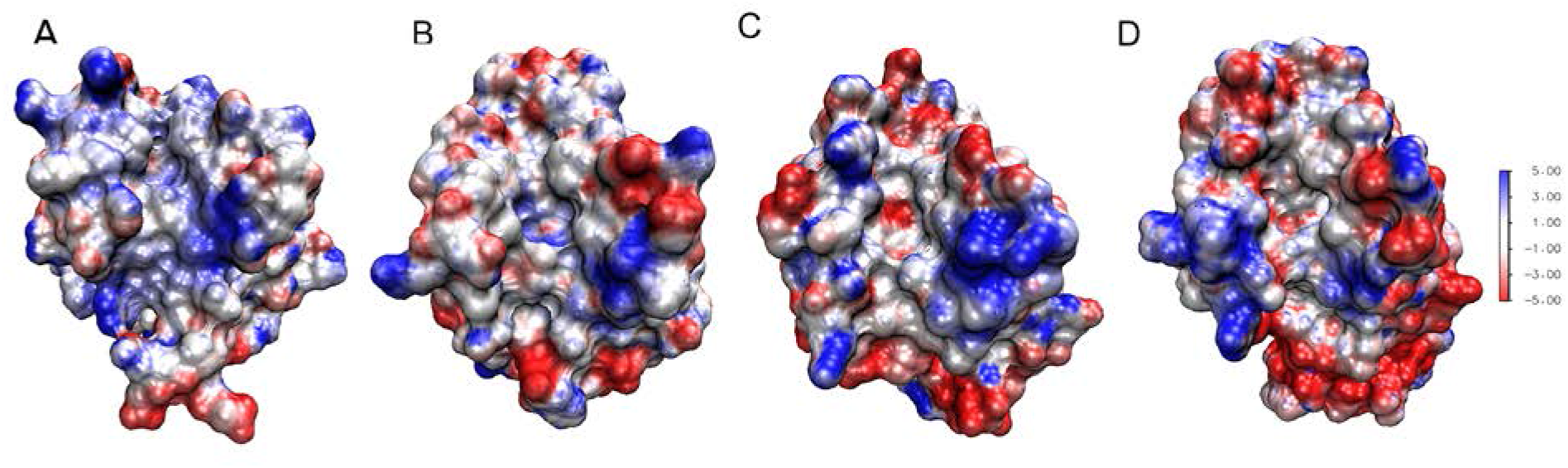
Electrostatic potential maps of (A) Mcl-1, (B) A179L, (C) BHRF1 and (D) Bcl-X_L_ calculated respectively using the experimentally determined structures with PDB IDs 2ROD, 5AU4, 2XPX and 2BZW. The electrostatic potentials were deduced by solving Poisson-Boltzmann equation utilizing Adaptive Poisson-Boltzmann Solver (APBS) ^60^ plugin of VMD software package ^61^. Red and blue colors in the diagrams correspond to negative and positive potentials respectively.

The BH3-like sequences show a similar trend while interacting with BHRF1 and are enriched with basic residues. MD simulations of selected BH3-like sequences demonstrate that interaction energies of basic residues are the most favorable with BHRF1. Electrostatic potential map of BHRF1 reveals several acidic patches (Figure 10C) indicating that the basic residues of BH3-like sequences can interact with these regions most favorably. However, unlike A179L, the acidic residues present in BH3-like sequences interact with BHRF1 with pronounced positive interaction energies (Figure 9). This is very well compensated by the favorable interaction energies of basic residues with BHRF1 which result in overall highly favorable Columbic interaction energies between the BH3-like sequences and BHRF1.

We have shown electrostatic potential maps of A179L and BHRF1 along with the bound BH3 peptides as representative examples (Figure 11). When A179L interacts with Bid wild-type BH3 peptide and Bad-5 the charged residues in both peptides result in favorable interactions (Figure 11A and 11B). This is clear from the interactions of acidic and basic residues of the peptides with the complementary regions in the protein. However, in the case of BHRF1, the acidic residues of Bak and Bak-c can be seen proximal to the negatively charged regions of BHRF1 especially towards the C-terminal segment of the peptide (Figure 11C and 11D). On the other hand, the basic residues of the peptides can be seen interacting with the acidic patches of the protein. This could be considered as one of the major differences between A179L and BHRF1. The other major distinguishing feature is the differences in the peptide:protein interaction energies between the wild-type BH3 peptides and the BH3-like sequences derived from them. In the case of BHRF1, this difference is significantly larger than that found for A179L.

**Figure 11:**
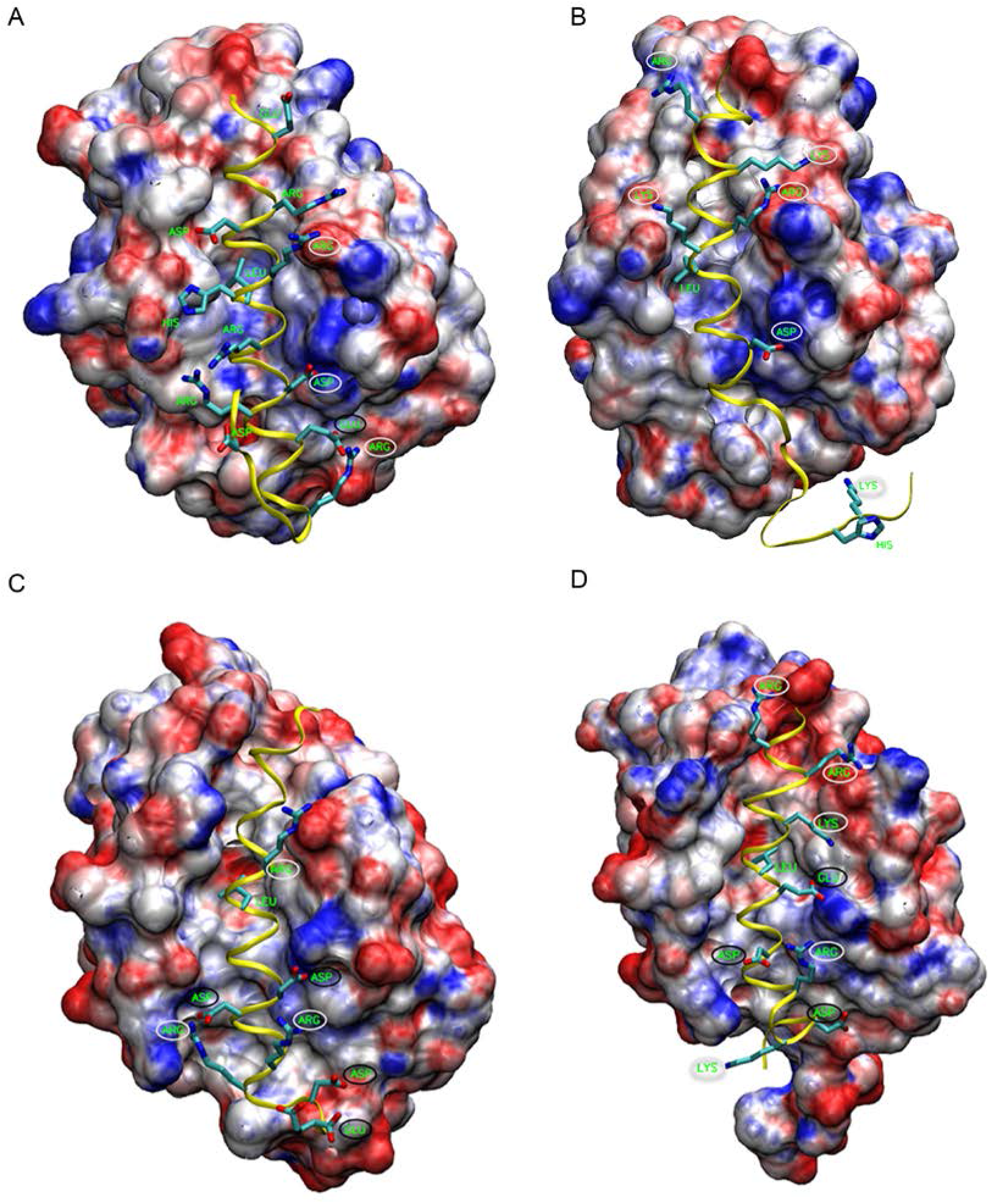
Electrostatic potential maps calculated for the MD simulated structures of A179 when it was bound to (A) wild-type Bid BH3 peptide and (B) Bid-based Bid-5 BH3-like peptide. Electrostatic potentials of MD simulated BHRF1 structures when the protein was in complex with (C) Bak wild-type BH3 peptide and (D) Bak-c, a BH3-like peptide derived from the wildtype Bak BH3 peptide. The MD simulated structures were saved at the end of 500 ns production runs. For details of electrostatic calculations, see Figure 10. Charged residues that are involved in favorable and unfavorable interactions are circled in yellow and black colors respectively.

## Discussion

The computational approach used in this study helped to design BH3-mimetic peptides that can bind and interact favorably with two viral Bcl-2 homologs A179L and BHRF1. These BH3-mimetic peptides are rich in basic residues and are involved in highly favorable long-range electrostatic interactions with the respective proteins. In a previous study, we carried out similar investigation on two mammalian anti-apoptotic Bcl-2 proteins, Mcl-1 and Bcl-X_L_^52^. A comparison of BH3-mimetic peptide sequences that bind to vBcl-2s and mammalian anti-apoptotic Bcl-2 proteins will help us to understand the major differences between these homologs and this knowledge is likely to provide an explanation why the inhibitors developed for mammalian pro-survival Bcl-2 proteins do not have the same effect in vBcl-2 homologs.

BH3-like sequences interacting favorably with Mcl-1 and Bcl-X_L_ showed distinct and specific preference for charged residues. BH3-like peptides enriched with acidic residues exhibited highly favorable interactions with Mcl-1 while the peptides with predominantly basic residues interacted favorably with Bcl-X_L_. Cell viability and cell proliferation studies on selected BH3-like peptides confirmed our prediction that these peptides indeed bind selectively to Mcl-1 or Bcl-X_L_ ^52^. Since the entropic component of ligand binding to hydrophobic grooves of the proteins is unfavorable ^62^, the free-energy of binding of BH3-peptides to anti-apoptotic proteins is largely dependent on enthalphic contribution. Thus the interaction energy between the peptide and the protein is an indicator of binding strength and affinity. Previous studies by Bond and coworkers also considered only the enthalphic component while studying the specificities and affinities of Bcl-2 family members ^63^. Hence, we extended our earlier approach to two vBcl-2 homologs, A179L and BHRF1. Both the vBcl-2 homologs considered in this study displayed preferences for BH3-like sequences with basic residues. Comparison of electrostatic potential of all the four proteins reveals that Mcl-1 has mostly basic patch around the BH3-binding pocket (Figure 10A). However in A179L, additional regions of negative electrostatic potential are found (Figure 10B) and the acidic patch has further expanded in BHRF1 (Figure 10C). Bcl-X_L_ seems to have the most acidic patch in the region where BH3 peptide binds (Figure 10D). This explains the preference of charged residues in BH3-like sequences that can bind to the vBcl-2 homologs A179L and BHRF1. While BH3-like sequences enriched in acidic residues bind favorably with Mcl-1, both acidic and basic residues interact favorably with A179L. Both BHRF1 and Bcl-X_L_ prefers to bind BH3-like sequences which are rich in basic residues. We have also argued that the presence of charged residues in the hydrophilic face of BH3 amphipathic helix can influence the binding of BH3 peptides as the electrostatic interactions are long range in nature. As a result, the so called non-hotspot residues seem to play a significant role in the affinity of the BH3-mimetic peptides. Although the four conserved hydrophobic residues in the BH3 region can help in binding the hydrophobic groove of the pro-survival proteins, other residues from the peptide can exert significant influence while interacting with the residues of the protein. Our current study along with our earlier observation that the long-range electrostatic interactions can determine the affinity of BH3-peptides in binding to anti-apoptotic Bcl-2 proteins have to be incorporated while designing inhibitors to both mammalian and vBcl-2 homologs.

## Conclusions

Although the Bcl-2 homologs in viruses are distantly related to the mammalian anti-apoptotic Bcl-2 proteins with strikingly similar three-dimensional fold, they have emerged attractive drug targets that will have the potential to prevent viral infection. African swine flu virus causes one of the most dangerous and fatal infectious diseases in domestic pigs and hence molecules that can inhibit viral infection can help economically many affected countries. Epstein-Barr virus has been implicated in human cancers including Hodgkin’s and Burkitt’s lymphoma. Structures of vBcl-2 homologs of both these viruses, A179L and BHRF1, adopt a typical Bcl-2 helical fold. With few studies available that focus on developing inhibitors to target A179L and BHRF1, the current study has used a computational approach to design peptides that can bind to the hydrophobic groove of these vBcl-2s. The designed peptide sequences selected from a library of BH3-like sequences have retained the original four conserved hydrophobic residues and a conserved Asp residue. The peptides have helical propensities significantly higher than the wild-type pro-apoptotic BH3 peptides. The unbiased selection of these peptides from the histogram of peptide:protein interaction energies revealed that the selected peptides are rich in basic residues. While A179L interacted favorably with both acidic and basic residues of the selected BH3-like peptides, the BHRF1 highly preferred peptides with basic residues and its interaction with acidic residues was clearly unfavorable. Analysis of electrostatic potential energy maps helped to understand the differences between mammalian anti-apoptotic Bcl-2 proteins and vBcl-2s. The Columbic interactions between the BH3-like peptides and the proteins predominantly contribute to the protein:peptide interaction energy. This is true for both the mammalian pro-survival Bcl-2 proteins and vBcl-2s. This approach can be extended to other vBcl-2s and the conclusions reached in this study can be incorporated in designing inhibitors that can specifically bind to A179L or BHRF1.

## Materials and Methods

### Generation of BH3-like sequences and modeling the vBcl2-BH3 peptide complex structures

We have adopted the same approach which we used to design BH3-mimetic peptides for the mammalian anti-apoptotic proteins Mcl-1 and Bcl-X_L_ in our earlier study ^52^. Briefly, a library of BH3-like peptides was generated from the selected wild-type pro-apoptotic BH3 peptide. In this step, the conserved four hydrophobic residues and the Asp residue were retained in their original positions and all other positions were substituted randomly without any bias. We considered the wild-type BH3 regions of eight pro-apoptotic proteins, namely, Puma, Noxa, Bmf, Bim, Bik, Bid, Bak and Bad and their BH3-domain sequences are shown in Figure 1. We employed “Random Protein Regions” module of Sequence Manipulation suite web server (http://www.bioinformatics./sms2/random_protein_regions.html) ^64^ for this purpose. For each pro-apoptotic BH3 peptide, we generated 1000 randomized BH3-like sequences.

The randomly generated BH3-like sequences were used to model complex structures using the experimentally determined BHRF1 or A179L structures. For all wild-type pro-apoptotic BH3 peptides for which complex structures are yet to be determined with A179L and for all randomly generated BH3-like sequences, the experimentally determined A179L:Bid complex structure (PDB ID: 5UA4) ^29^ was used as template structure. Structures of BHRF1 in complex with Bim (PDB ID: 2WH6) and Bak (PDB ID: 2XPX) are available in PDB ^20^. However, examination of these structures reveals that there are missing regions in both the structures and hence we used both as template structures while modeling wild-type pro-apoptotic BH3 peptides or BH3-like sequences generated randomly. We used Modeller 9.14 ^65–67^ to model the complex structures and the protocol used to model the complex structures is the same as described previously^52^. For each BH3-like sequence, we used this modeling procedure to generate five best models which was further energy minimized in a water box using GROMACS 4.5.5 software suite ^68^. Each modeled protein-peptide complex structure was solvated using SPC/E water model ^69^ and OPLS/AA force-field ^70^ was used in the minimization protocol. Short- and long-range interactions were evaluated for the non-bonded pairs that were within a cut-off of 10 Å and within 10-25 Å respectively. Among the five selected models, the model with minimum potential energy was selected to calculate the interaction energy between the peptide and protein as given in the following equation.

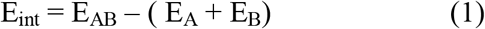

Where E_ab_, E_a_ and E_B_ represent the potential energies of the complex structure, the protein A179L/BHRF1 and the wild-type BH3/BH3-like peptide respectively. Interaction energies were calculated between the wild-type BH3 or the BH3-like sequences and the protein. In total we modeled 16000 structures (1000 BH3-like sequences generated each for the eight wild-type BH3 sequence and the two sets of 8000 sequences thus generated were used to build A179L and BHRF1 complex structures). Histograms of interaction energies for A179L and BHRF1 complex structures were plotted for BH3-like sequences derived based on each of the eight wild-type BH3 sequences. We also determined the helicity of each BH3-like sequence using Agadir webserver at 278K, pH 7 and 0.1 M ionic strength ^56–58^. Amphipathic nature of each BH3-like sequence was determined by quantifying the mean helical hydrophobic moment using HELIQUEST webserver^59^

### Molecular Dynamic simulation of selected complex structures

We performed molecular dynamics simulations of selected models of A179L and BHRF1 structures in complex with BH3-like sequences. Simulations were carried out following the same^52^ protocol used in our earlier studies. In each case, the complex structure was first solvated using SPC/E water model ^69^ in a cubic box and the minimum distance between the complex and the wall of the box was at least 13 Â. The systems were neutralized by adding the required number of counter-ions and then energy minimized. OPLS all-atom force-field^70^ was used and the simulations were performed using GROMACS 4.5 ^68^. Equilibration step consisted of 1 ns simulation in NVT ensemble with positional restraints (force constant 2000 kJ mol^-1^ nm^-1^) on all heavy atoms. This was followed by 1.8 ns simulation in NPT ensemble in which the positional restraints were gradually removed on all atoms except the backbone atoms. Restraints on backbone atoms were gradually removed over a period of another 2 ns simulation in NPT ensemble to ensure that the modeled BH3 peptide helix is properly equilibrated. After the equilibration, each system was simulated for a period of 500 ns. The system was maintained at the reference temperature 300 K. All other details were same as given in Reddy et al. ^52^. Summary of all the systems simulated is provided in Table S1. A total of 7 p,s simulation was carried out for all the systems. The coordinates were saved every 10 ps and 50,000 MD-simulated structures were used for further analysis for each system. Helix stability of BH3-like peptides, interaction energies between peptide and protein and contribution of acidic and basic residues for the protein-peptide interaction energies were analyzed in MD simulations. Interaction energy between the protein and the peptide for each MD simulated structure was calculated using the Eq. (1). A twin-range cut-off of 10 to 25 Â was used for the purpose of calculating the interaction energy between the peptide and protein for all the MD-simulated structures.

## Acknowledgements

We gratefully acknowledge the High Performance Computing Facility at IIT-Kanpur. RS is Pradeep Sindhu Chair Professor. CNR thanks BINC Fellowship from the Department of Biotechnology (DBT), Government of India.

