## Supporting Information for "Designing BH3-mimetic Peptide Inhibitors for the Viral Bcl-2 Homologs A179L and BHRF1: Importance of long-range electrostatic interactions"

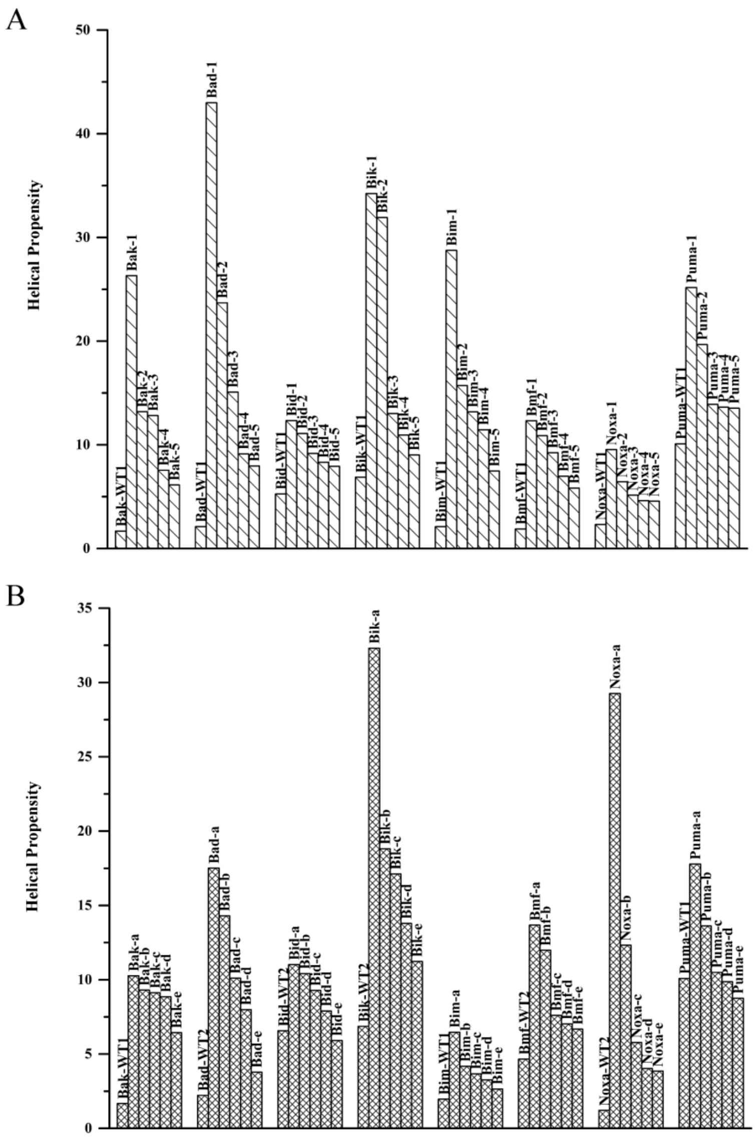


**Figure S1**: Helical propensities of wild-type BH3 and selected BH3-like sequences that bind to (A) A179L or (B) BHRF1 with highly favorable interaction energies. Helical nature of all wild-type BH3 and BH3-like sequences were determined using AGADIR web server (http://agadir.crg.es).

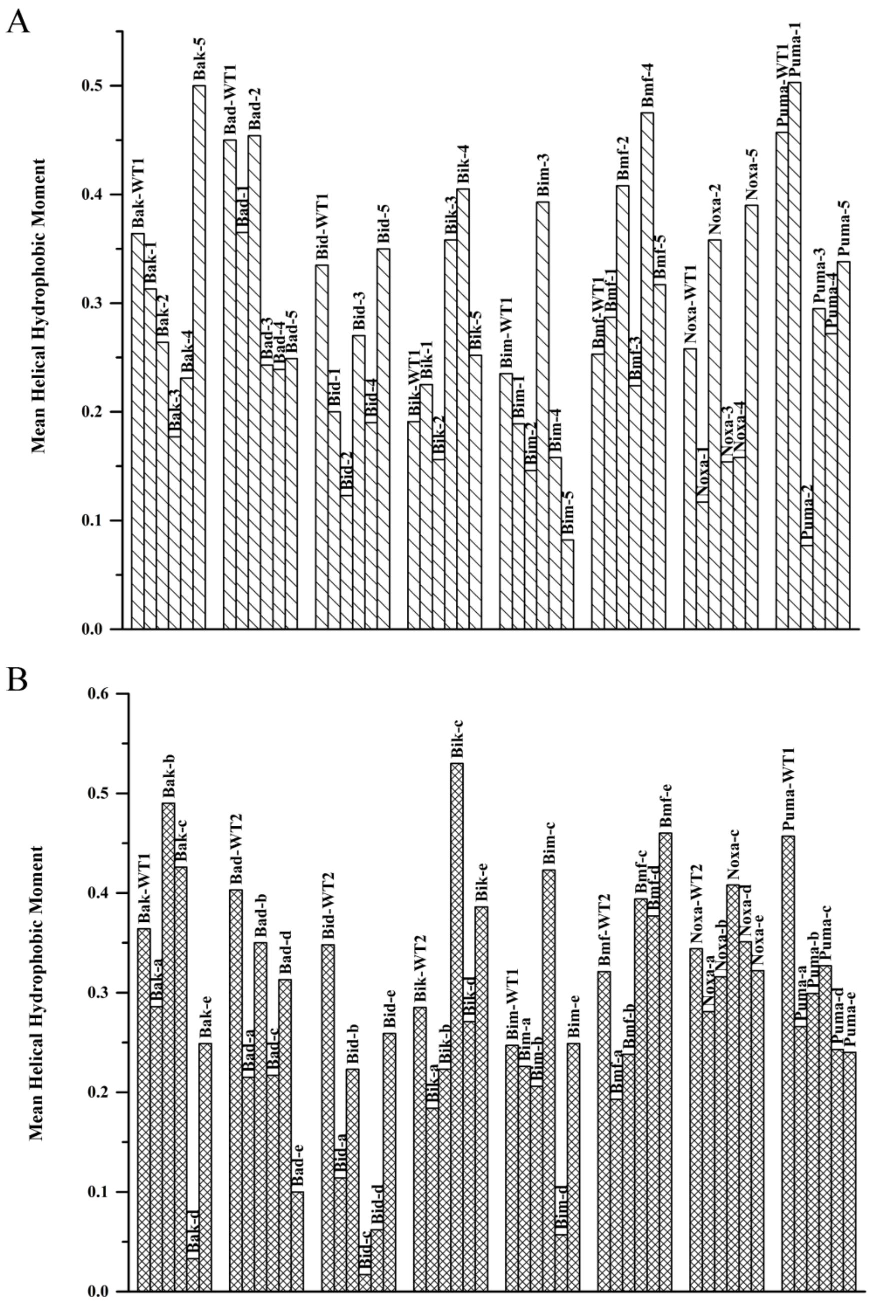


**Figure S2**: Mean helical hydrophobic moment of wild-type BH3 and selected BH3-like sequences that bind to (A) A179L or (B) BHRF1 with highly favorable interaction energies. Mean helical hydrophobic moment was calculated to HELIQUEST web server (http://heliquest.ipmc.cnrs.fr).

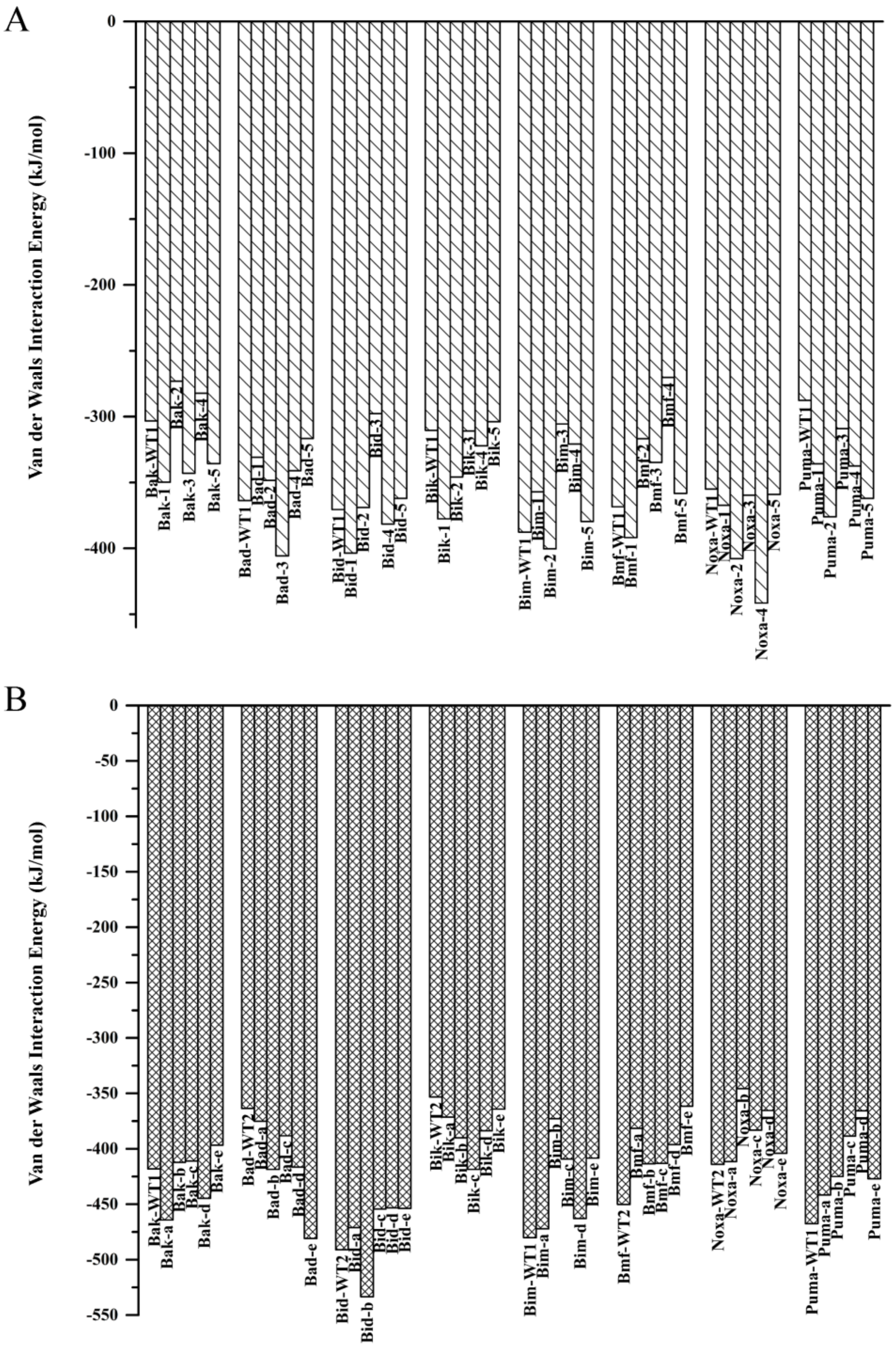


**Figure S3**: van der Waals component of interaction energies between BH3 peptides (wild-type and BH3-like) and (A) A179L and (B) BHRF1 vBcl-2 homologs.

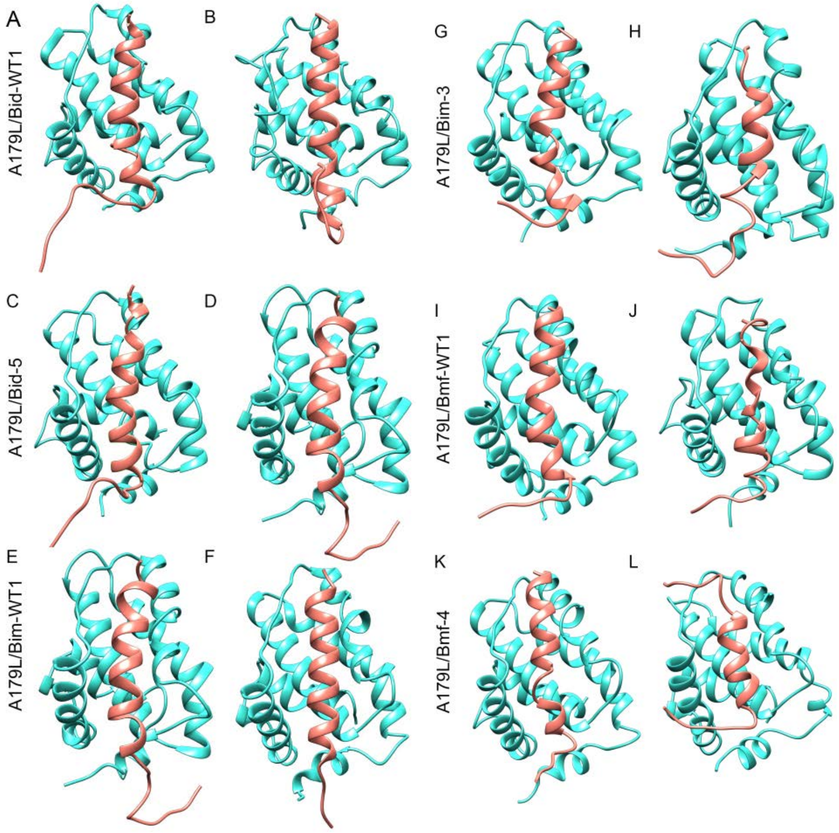


**Figure S4**: The vBcl-2 homolog A179L in complex with (A,B) Bid-WT1, (C,D) Bid-5, (E,F) Bim-WT1, (G,H) Bim-3, (I,J) Bmf-WT1 and (K,L) Bmf-4. The initial structures for MD simulations (A,C,E,G,I,K) and the structures saved at the end of 500 ns simulations (B,D,F,H,J,L) are shown. For summary of all simulations, see Table S1.

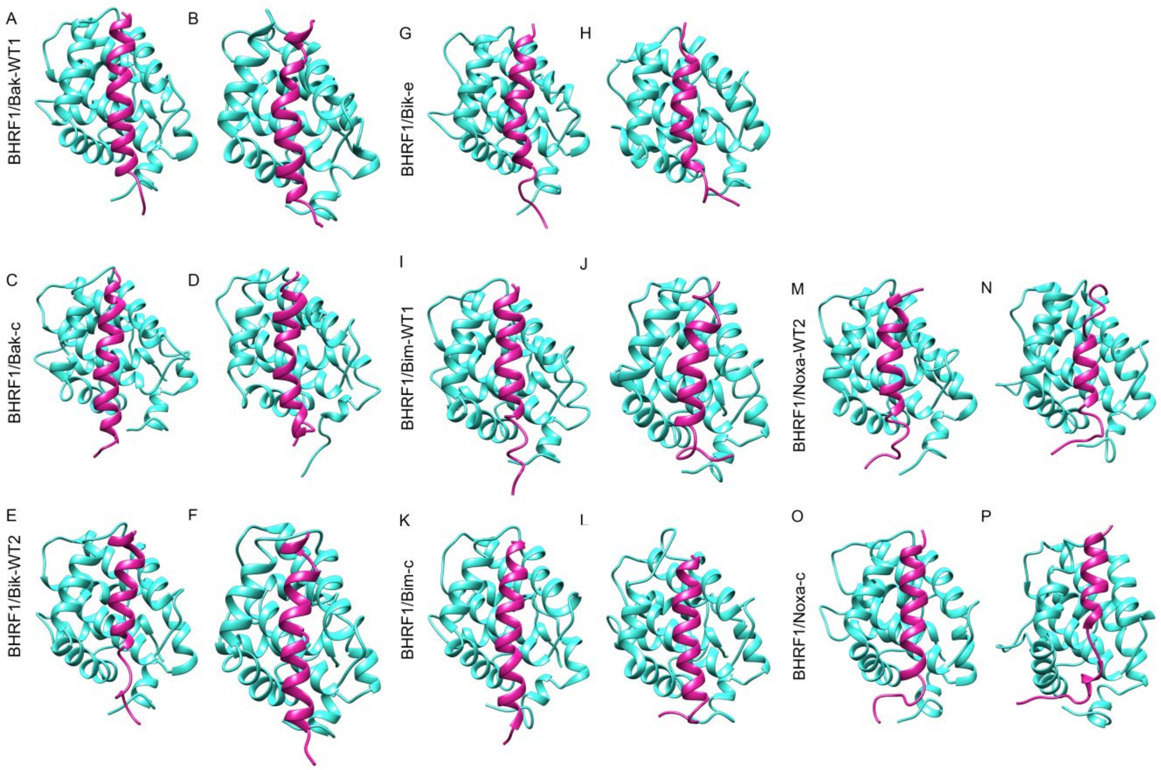


**Figure S5**: The vBcl-2 homolog BHRF1 in complex with (A,B) Bak-WT1, (C,D) Bak-c, (E,F) Bik-WT2, (G,H) Bim-e, (I,J) Bim-WT1, (K,L) Bim-c, (M,N) Noxa-WT2 and (O,P) Noxa-c. The initial structures for MD simulations (A,C,E,G,I,K,M,O) and the structures saved at the end of 500 ns simulations (B,D,F,H,J,L,N,P) are shown. For summary of all simulations, see Table S1.

**Table S1**: Summary of molecular dynamics simulations of viral Bcl-2 homologs A179L and BHRF1 in complex with pro-apoptotic wild-type BH3 peptides or BH3-like peptides.

| S. No. | Protein:peptide complex system[a] | Production run |
|---|---|---|
| 1. | A179L:Bid-WT1 | 500 ns |
| 2. | A179L:Bid-5 | 500 ns |
| 3. | A179L:Bim-WT1 | 500 ns |
| 4. | A179L:Bim-3 | 500 ns |
| 5. | A179L:Bmf-WT1 | 500 ns |
| 6. | A179L:Bmf-4 | 500 ns |
| 7. | BHRF1:Bak-WT1 | 500 ns |
| 8. | BHRF1:Bak-c | 500 ns |
| 9. | BHRF1:Bik-WT2 | 500 ns |
| 10. | BHRF1:Bik-e | 500 ns |
| 11. | BHRF1:Bim-WT1 | 500 ns |
| 12. | BHRF1:Bim-c | 500 ns |
| 13. | BHRF1:Noxa-WT2 | 500 ns |
| 14. | BHRF1:Noxa-c | 500 ns |

[a]Sequences of BH3 wild-type peptides and BH3-like sequences can be found in Figure 1, Figure 4 and Figure 6.
